# Defining p53 pioneering capabilities with competitive nucleosome binding assays

**DOI:** 10.1101/327635

**Authors:** Xinyang Yu, Michael J. Buck

## Abstract

Accurate gene expression requires the targeting of transcription factors (TFs) to regulatory sequences often occluded within nucleosomes. The ability to target a transcription factor binding site (TFBS) within a nucleosome has been the defining characteristic for a special class of TFs known as pioneer factors. Recent studies suggest p53 functions as a pioneer factor that can target its TFBS within nucleosomes, but it remains unclear how p53 binds to nucleosomal DNA. To comprehensively examine p53 nucleosome binding we competitively bound p53 to multiple *in vitro* formed nucleosomes containing a high or low-affinity p53 TFBS located at differing translational and rotational positions within the nucleosome. Stable p53-nucleosome complexes were isolated and quantified using next generation sequencing. Our results demonstrate p53 binding is limited to nucleosome edges with significant binding inhibition occurring within 50-bp of the nucleosome dyad. Binding site affinity only affects p53 binding for TFBS located outside the nucleosome core at the same nucleosomal positions. Furthermore, p53 has strong non-specific nucleosome binding facilitating its interaction with chromatin. Our *in vitro* findings were confirmed by examining p53 induced binding in a cell line model, showing induced binding at nucleosome edges flanked by a nucleosome free region. Overall, our results suggest that the pioneering capabilities of p53 are driven by non-specific nucleosome binding with specific binding at nucleosome edges.

## INTRODUCTION

Appropriate gene expression is vital for the success of countless cellular processes including growth, development, and metabolism. Among the factors known to regulate gene expression is a highly characterized subset of proteins known as transcription factors (TFs). TFs regulate genes by binding to specific DNA sequences in the genomic vicinity of each regulated gene and inducing a change in expression. TFs recognize degenerate sites as transcription factor binding sites (TFBS) that appear thousands of times across a given eukaryotic genome. *In vivo,* most TFBSs are never targeted, likely due to the fact that they are inaccessible due to binding by nucleosomes.

Nucleosomes are the primary unit of chromatin structure comprised of a 147bp section of DNA wrapped around a histone protein core with neighboring nucleosomes separated by accessible linker DNA (Kornberg and Lorch 1999; Jiang and Pugh 2009). In theory, the steric hindrance of nucleosome-DNA interactions could inhibit TFs binding to all nucleosome-bound DNA. In reality, however, nucleosome inhibition of TFs binding is variable both across a genome and even within a single nucleosome (Buck and Lieb 2006). As DNA duplex molecule bends and twists around the histone octamer, one side of it directly contacts the histone surface and gets buried inside, while the other side is exposed to the solvent and accessible, with a concealed/exposed periodicity of 5-bp. This well-organized architecture maintains nucleosome stability via electrostatic interactions and hydrogen bonds between histone protein side chains and phosphate groups in the DNA backbone. As a result, this highly distorted and partially overlaid nucleosomal DNA cannot be accessed readily by TFs (Vermaak et al. 2003; Luger et al. 2012).

The location of TFBS within a nucleosome can significantly affect TF-nucleosome binding. TFBS can have various positions within a nucleosome core (known as translational setting), from near the edge to the center of the nucleosome dyad. For the glucocorticoid receptor a TFBS located near the edge is bound 4-fold better compared to an identical TFBS positioned 20-bp from the dyad (Li and Wrange 1993). Other TFs are inhibited by the translational settings by differing amounts from 2-100 fold (Vettese-Dadey et al. 1994; Blomquist et al. 1996; Angelov et al. 2004). These divergences in nucleosome inhibition of TF binding are likely driven by differences in how TFs recognize their binding sites on nucleosomes. The orientation of a TFBS on a nucleosome (known as “rotational setting”) also influences how nucleosomes inhibit TF binding. TFBSs located along a nucleosome surface can either face inward or outward, due to the twisting of DNA’s helical structure. For FoxA a TFBS located near the nucleosome dyad at a specific rotational setting is significantly bound in *in vitro* binding assays while a rotational shifted TFBS located 5-bp away is not bound (Sekiya et al. 2009).

FoxA represents a special class of transcription factors known as pioneer factors. Pioneer factors were first described in 2002, as regulatory proteins capable of targeting DNA sequences even within compacted, closed chromatin, while other TFs cannot (Cirillo et al. 2002). Among all known TFs, only a few have been characterized as pioneer TFs (Iwafuchi-Doi and Zaret 2014). Recent studies suggest that p53 functions as a pioneer factor at some of its binding sites (Sammons et al. 2015). Sammon et al (2015), showed that activated p53 bound to 4,416 new bindings sites, 44% of which reside within inactive (H3K4me1-and H3K4me3-) and inaccessible chromatin (Sammons et al. 2015). At the p21 gene p53 binds at a higher affinity to its binding site within chromatin compared to naked DNA (Espinosa and Emerson 2001). Furthermore, throughout the human genome p53 TFBS occur in regions with strong nucleosome positioning sequences (Lidor Nili et al. 2010). These findings suggest that p53 functions as a pioneer factor that can target its binding sites within nucleosomes.

p53 is a DNA binding transcription factor that acts as a tumor suppressor and with its DNA-binding domain being frequently mutated during cancer (Rivlin et al. 2011). p53 integrates multiple stress-induced signals and acts as a transcriptional regulator for a wide variety of genes involved in DNA repair, cell-cycle arrest, and apoptosis. p53’s ability to regulate these transcriptional responses requires it to specifically target and bind its binding sites throughout the genome. p53’s recognition of naked DNA has been extensively studied and binds to DNA as a homo-tetramer recognizing two decamers of RRR<u>CWWG</u>YYY (el-Deiry et al. 1992), each decamer is called a ‘half site’ with underlined 4 nucleotides CWWG as the core, usually one full site consists of two half sites. p53 binding to nucleosomes has been previously tested with conflicting findings. Laptenko *et al.* showed that a p53 TFBS located near the dyad had un-detectable binding, while sites near the nucleosome edge are bound (Laptenko et al. 2011). Sahu *et al.* showed that p53 can bind to sites near the nucleosome dyad when in the appropriate rotational position (Sahu et al. 2010). Therefore, to address these conflicting results we developed a quantitative and competitive binding assay allowing the direct comparison of multiple nucleosome sequences in a single binding assay.

To comprehensively examine p53 nucleosome binding, we competitively bound p53 to multiple nucleosomes containing a p53 binding site (p53BS) located at differing translational and rotational positions within the nucleosome. Differences in binding affinity were then quantified using next generation sequencing. Our results demonstrate p53 sequence specific binding is limited to nucleosome edges with significant binding inhibition occurring within 50-bp of the nucleosome dyad. Our *in vitro* findings were confirmed by examining p53 induced binding in a cell line model, showing induced binding at nucleosome edges flanked by a nucleosome free region. In addition to sequence specific binding p53 binds relatively strongly in a sequence-independent manner to nucleosomes. Overall, our results suggest that the pioneering capabilities of p53 are driven by non-specific nucleosome binding with specific binding at nucleosome edges.

## RESULTS

### Nucleosome design and formation

To determine the specificity of p53 to nucleosomal DNA we developed competitive nucleosome binding assay (Fig. 1A). Starting with the Widom 601 nucleosome positioning sequence, 14 templates were designed and compared to non-specific binding to 2 control sequences (Fig. 1B). The Widom 601 nucleosome positioning sequence produces highly stable and optimally positioned nucleosomes (Lowary and Widom 1998; Anderson and Widom 2000). With increasing distance to the dyad axis, three translational settings were tested – dyad (superhelix location (SHL) 0, 0.5), intermediate (SHL 4, 4.5), and edge (SHL 6.5, 7) (Fig. 1B). At each translational position two rotational settings were tested by shifting the p53BS 5-bp to the right. TFBS accessibility was determined by modeling the TFBS position onto the nucleosome crystal structure formed from the Widom 601 sequence (Makde et al. 2010). In addition, a site in the linker region (SHL 8), which is outside the nucleosome core, was examined as well.

**Figure 1.**
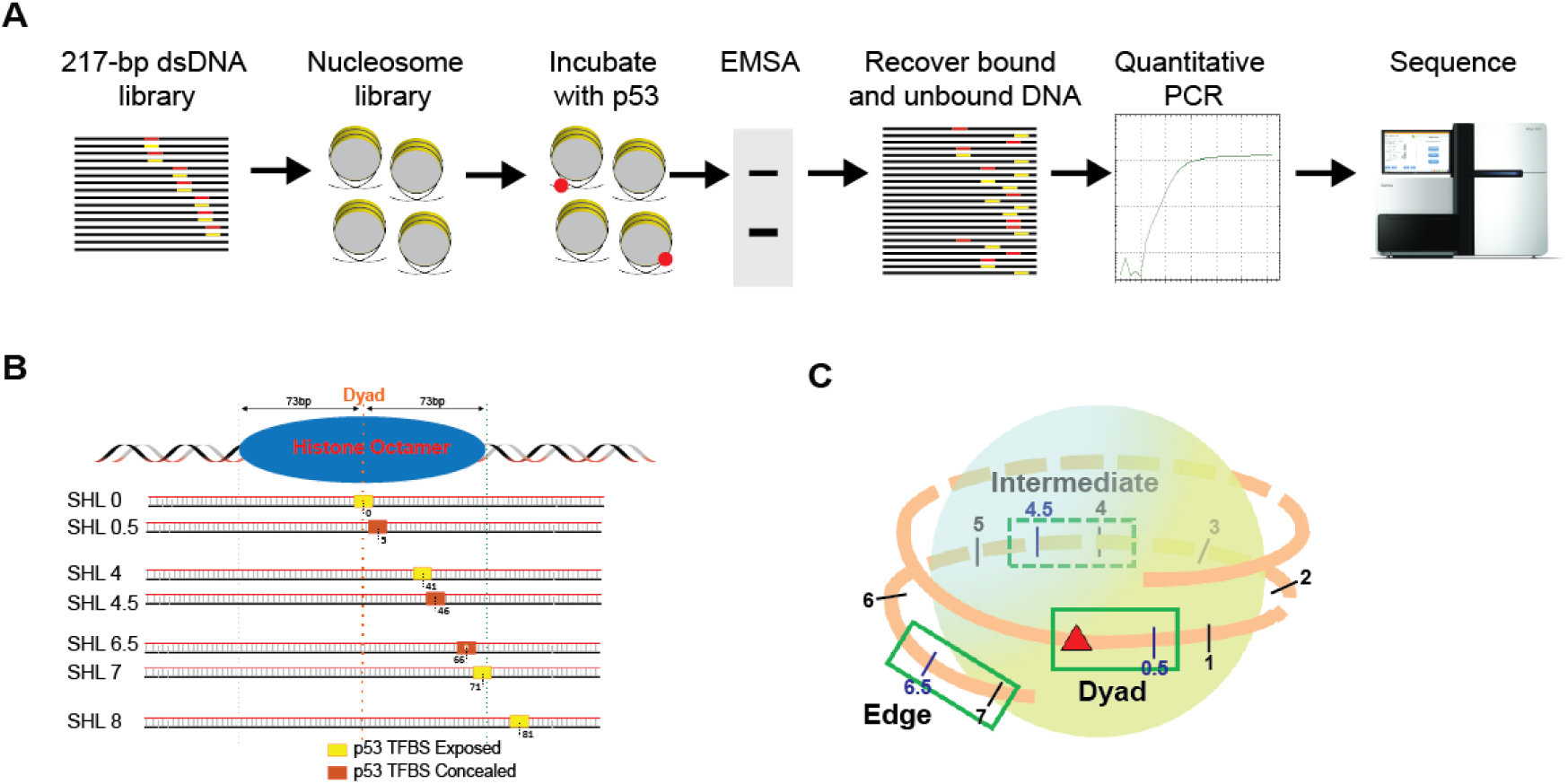
Design of nucleosome templates with p53BS. **(A)** 217-bp dsDNA library was designed containing p53BS in various nucleosomal positions. Nucleosomes were formed, and purified generating a nucleosome library that was incubated with increasing amounts of p53 protein. p53-nucleosome complexes were separated by EMSA and the bound and un-bound DNA was recovered, quantified by qPCR, and sequenced. **(B)** A 20-bp-long p53BS (low-and high-affinity) was placed at different positions with 0-bp, 5-bp, 41-bp, 46-bp, 66-bp, 71-bp, 81-bp away from the dyad. The superhelix location (SHL) is designated for each nucleosome sequence. **(C)** Within the nucleosome 3 different positions (dyad, intermediate, and edge) were chosen with increasing distance to nucleosome dyad.

We selected two p53BS, the first is from its *in vivo* target p21 promoter with relative low affinity: 5’-AGA<u>CTGG</u>GCATGT<u>CTGG</u>GCA-3’ (Westfall et al. 2003), the other one is a modified sequence with high p53 binding affinity: 5’-GGG<u>CATG</u>TCCGGG<u>CATG</u>TCC-3’ (Veprintsev and Fersht 2008; Noureddine et al. 2009). The core sequences CTGG and CATG, for the low and high-affinity sites respectively, directly interact with the p53 protein. The functional p53 full site is 20-bp composed of two 10-bp p53BS half-sites. This 10-bp half-site length allows both half-sites to remain in-phase on nucleosomal DNA. The two p53BS were then separately added to Widom 601 DNA by replacing the base pairs at the selected locations. Consequently, we obtained 14 different nucleosomal templates having p53BS with increasing distance to the nucleosome dyad, and being placed in either exposed or concealed orientation (Fig. 1C).

The 217-bp nucleosome sequences for these experiments were obtained as double stranded DNA fragments. Each nucleosome sequence was amplified and then pooled at equal molar concentrations. Nucleosomes were assembled using salt gradient dialysis on all nucleosome sequences simultaneously (Hayes and Lee 1997). Nucleosome positioning was then examined by native PAGE (Fig 2A). Native PAGE has been shown to be an optimal quality control test to assess *in vitro* reconstituted mono-nucleosomes because of its high sensitivity to shape and charge. This traditional method successfully differentiates different nucleosome positions on the same DNA sequence (Dyer et al. 2004; van Vugt et al. 2009; Muthurajan et al. 2016; Silberhorn et al. 2016). Our EMSA results show a single nucleosome band, confirming the correct positioning for all 16 nucleosome sequences (Fig. 2A).

**Figure 2.**
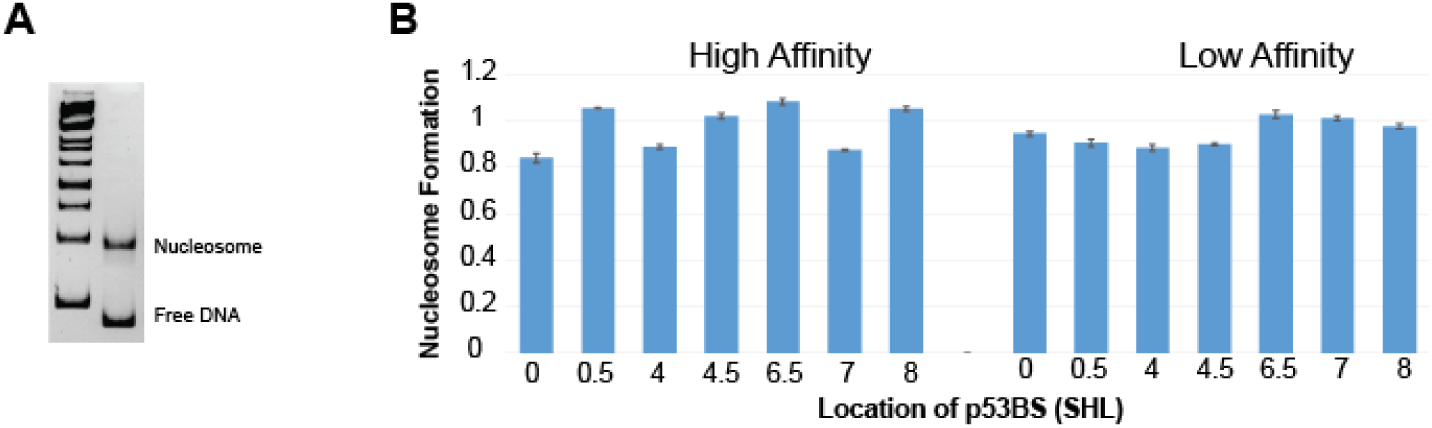
Nucleosomes form efficiently at a single position. **(A)** EMSA separating nucleosomes from free DNA before nucleosome purification. Only a single nucleosome band is present indicating a single nucleosome position on all templates after *in vitro* nucleosome reconstitution. **(B)** Nucleosome formation efficiency is determined by counting the frequency of each sequence within the nucleosomes compared to the total frequency of each sequence in both the free DNA and nucleosomes relative to formation for control 601 sequence (see Eq. 1). Nucleosome formation was determined for every single nucleosome template used in the experiment. Error bars are standard error of the mean for 4 replicate experiments.

Since we are modifying the Widom 601 nucleosome positioning sequence by inserting p53BS, we validated nucleosome formation efficiency for each sequence as compared to the un-modified Widom 601 sequence. After nucleosome assembly and gel shift assay, the nucleosomal DNA and free DNA were gel-extracted and sequenced (Fig. 2A). By comparing the number of reads for each template sequence in the nucleosome band with the number of reads for 601 control sequence we can determine the relative nucleosome formation efficiency (see Eq. 1). Nucleosome formation efficiency for each template was extremely consistent across replicates with only small differences among nucleosome templates, less than 20% (Fig. 2B).

### p53 is occluded from binding nucleosome core region

To get a comprehensive idea of how p53 binds to nucleosomes, we combined the traditional TF-nucleosome binding assay with high throughput sequencing, so that we can analyze multiple positions on the nucleosome simultaneously with one binding reaction. We added p53 protein to 0.5 pmol purified nucleosome with increasing amount of p53 (0 to 2 pmol, 0 to 286 nM). After a short incubation, the binding reactions were separated on a native polyacrylamide gel to detect the p53-nucleosome complex (Fig. 3A). The first lane contained only nucleosomes and was used to measure background and input levels for each experimental replicate. As the concentration of p53 increased, the first supershift band intensity increased and higher order bands appeared. It is also noteworthy that as p53 amount increased, the intensity of nucleosome-only band significantly decreased, and even completely disappeared when p53 amount reached 286 nM, indicating a total binding and shifting of all nucleosomes even including the 12.5% (2 out of 16) no-p53BS-containing control nucleosomes. To determine the makeup of the shifted complex we performed a modified western blot from the EMSA gel (see Methods). This result for p53 and Histone H4 both confirm a stable ternary complex composed of p53 and nucleosome (Fig. S1).

**Figure 3.**
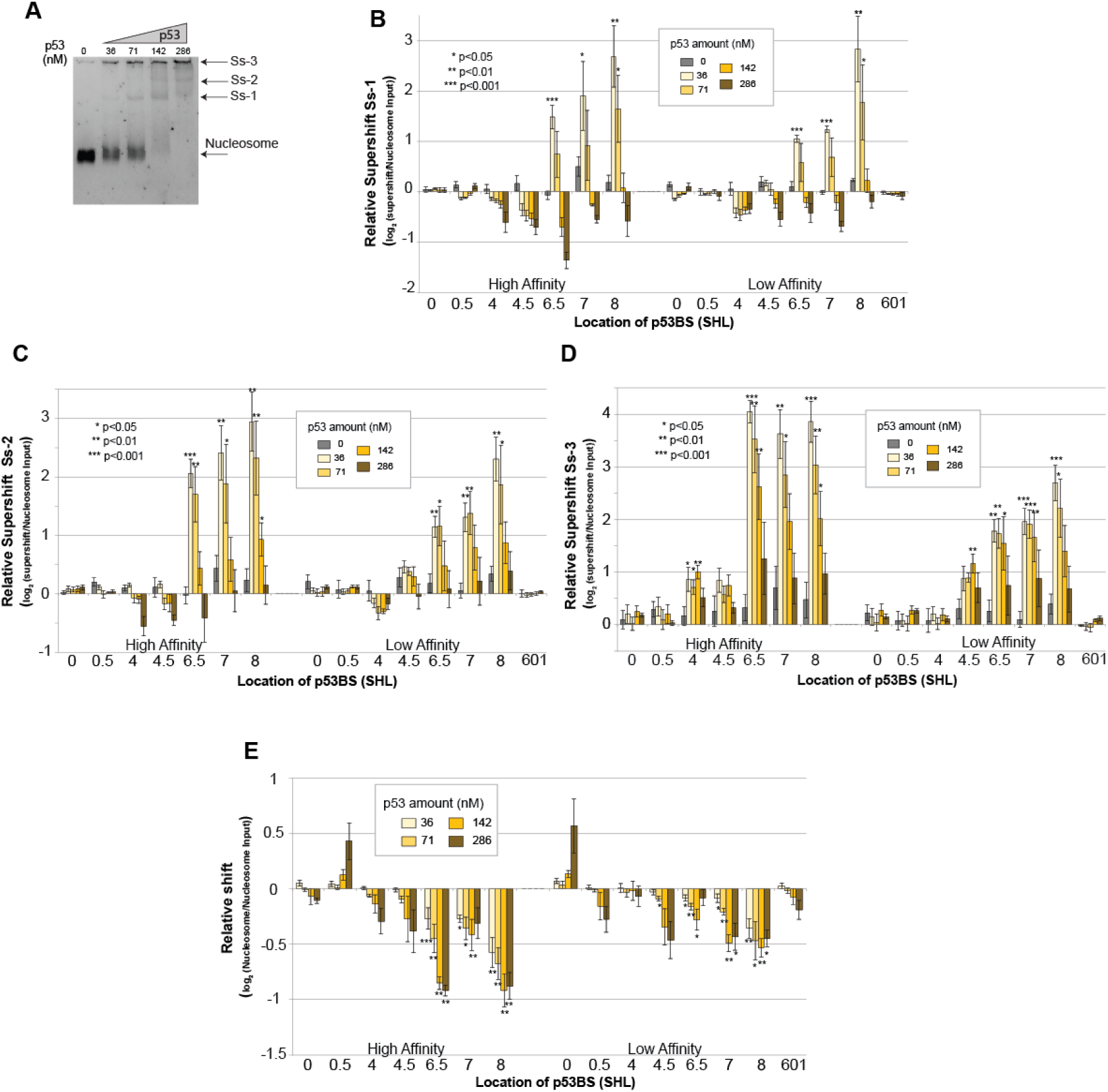
p53 is occluded from binding the nucleosome core region. **(A)** Nucleosomes containing 16 different sequences were bound to increasing amounts of p53 protein and separated by native PAGE. Lanes contain the following:1– 0.5 pmol inputed nucleosomes, 2-5 contain 0.5 pmol of nucleosomes with 36, 71, 142, or 286 nM of p53 (0.25, 0.5, 1, or 2 pmol). Nucleosome and the major supershift bands are indicated (Ss-1, Ss-2, Ss-3). **(B-D)** Relative supershift for each nucleosome is determined by counting the frequency of each sequence within the supershift and comparing it to non-specific binding to the 601 sequence. This value is then normalized to the input ratios of nucleosomes (see Eq. 3). Error bars are standard error of the mean; p-values are shown comparing each nucleosome sequence at a specific p53 concentration to background levels in the input lane. **(B)** First-supershift band (Ss-1) **(C)** Second-supershift band (Ss-2) **(D)** Third-supershift band (Ss-3) **(E)** Nucleosome sequences that are lost from the nucleosome band were quantified relative to the input nucleosomes. Strongly bound nucleosome will generate negative relative shifts.

To determine which nucleosome p53 preferentially binds, all shifted bands were gel extracted, DNA purified, quantified by qPCR, and sequenced. The sequencing results were then analyzed and mapped back to the original 16 nucleosome sequences. The resulting dataset was then analyzed by multiple methods all producing the same conclusions. In the first method raw sequence counts were analyzed by comparing each template sequence to the control non-specific sequence in the same binding experiment (see Eq. 2). By comparing each nucleosome sequence to the non-specific control sequences from the same lane, loading, PCR amplification, and next-generation sequencing are all internally controlled. This approach is performed on each shifted band independently (Fig. 3). Statistical significance for each template sequence can then be determined compared to background measurements from the p53-null lane. This analysis demonstrates that nucleosome sequences containing a p53 TFBS located outside of the nucleosome are bound first at the lowest concentrations. Sites located near the nucleosome edge are also bound at low concentrations, while the nucleosomes containing a p53 TFBS located near the dyad (+/-50-bp) are not specifically bound as compared to the control nucleosomes. At higher concentrations of p53, 142 and 286 nM, binding is no longer significantly different than non-specific binding to the control nucleosomes.

To confirm these result from the shifted bands we also examined the non-shifted nucleosome fragments (Fig 3E). In this analysis we examined the loss of DNA fragments relative to the starting amount (p53-null nucleosome band). Nucleosomes which are bound strongly by p53 will be shifted out of the nucleosome band and generate a negative relative shift. The advantage of this approach is that it allows the determination of binding regardless of p53/nucleosome complex structure and location of the supershift. Previous studies have shown that p53 oligomerizes into larger complexes (Stenger et al. 1992; Lee et al. 1994; Chene 2001; Kearns et al. 2016). These results confirm specific binding at nucleosome edges and linker. In this analysis we do not see a drop-off in binding at higher p53 concentrations relative to the control because once a specific nucleosome is shifted away from the nucleosome it will still be accurately counted as missing, regardless of where it has been shifted too. This differs from our examination of the supershift fragments because at higher concentrations of p53, higher/order complexes are created further shifting the nucleosome sequence and removing them from the supershift count.

The two approach above are unable to estimate the non-specific binding of p53 to nucleosomes, therefore to determine both specific and non-specific binding directly we performed a more in-depth analysis using the DNA amounts determined by qPCR. Briefly, after isolating the DNA from the native PAGE the amount of DNA was determined by qPCR using a standard curve generated from a control DNA fragment. This allows us to determine the absolute number of DNA molecules in a shifted or nucleosome band before amplification for NGS library generation. After sequencing we then convert the relative counts from the sequencing library to an absolute number (Eq 3.). These absolute counts were then used to determine the % nucleosome bound for each template across all p53 concentrations (Eq. 4). The K_D_ was then determined by non-linear regression (Heffler et al. 2012b). As seen with the previous methods, significant supershift occurs for templates with p53 TFBS at the edge or outside of the nucleosome (Fig. 4). In both high-affinity p53BS and low-affinity p53BS groups, it is clear that p53BS located near the nucleosome edges have the smallest K_D_ values. As the p53BS moves closer into the nucleosome dyad, K_D_ value gradually increases (Fig. 4C), and eventually show no significant difference from the control non-BS containing nucleosomes. Interestingly, p53BS affinity only matters when it is placed at nucleosome edges or in the linker, there is no significant difference between high-affinity p53BS and low-affinity p53BS when they are located within 50-bp of the nucleosome dyad (Fig. 4C). All analyses approaches consistently demonstrate that only p53BS located within the nucleosome edge are bound. Furthermore, binding site affinity only affects p53 binding for TFBS located at the same nucleosomal positions. When we look at the same site in the linker region, the high-affinity p53BS has a lower K_D_ than low-affinity p53BS, but when we compare the high-affinity p53BS in the edges (SHL 7) with the low-affinity BS in the linker (SHL 8), the low-affinity group displays smaller K_D_ values indicating higher binding affinity. This result shows that nucleosome translational position takes precedent over binding site affinity.

**Figure 4.**
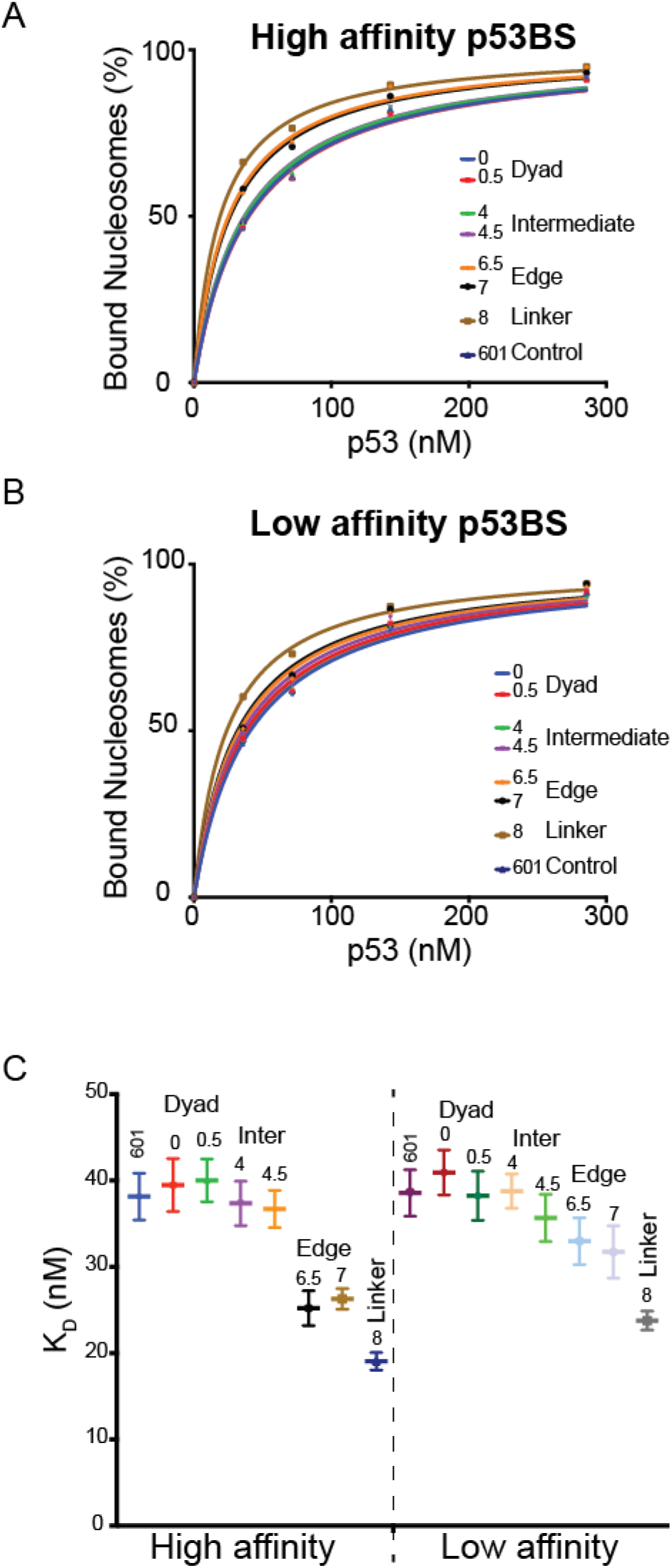
p53 has higher affinity to BS near nucleosome boundaries. **(A-C)** The sequence results for the nucleosome bands were used to determine the binding affinities, K_D_ with non-linear regression. **(A)** The percent of nucleosomes bound by p53 as p53 is titrated. The location of the high-affinity p53BS is indicated. **(B)** The percent of nucleosomes bound by p53 as p53 is titrated. The location of the low-affinity p53BS is indicated. **(C)** K_D_ values with standard errors were plotted for each nucleosome.

To our surprise p53 binds relatively strongly to all nucleosome fragments regardless of the presence or absence of a p53BS. At the highest concentrations of p53 all nucleosomes appear to shift regardless of the presence or absence of a p53 TFBS. In particular, there is a little over a 2-fold difference in kD between linker associated p53BS and the control nucleosomes. This 2-fold difference in sequence-dependent and sequence-independent binding affinity is similar to what was seen for FoxA pioneer transcription factor (Sekiya et al. 2009).

### p53’s preference to nucleosome boundaries is not driven by DNA sequence

To confirm the differences seen in our nucleosome binding experiments are not due the placement of the p53BS at various locations within 601 we performed a control binding experiment to the pool of 16 DNA sequences. In this assay, all 16 DNA templates were mixed at equal amount and then added to p53 protein. The DNA band did not disappear even as we increased the p53 protein amount to as high as 4 pmol (0.57 µM) for binding with 0.5 pmol DNA (Fig. 5A). The p53-DNA supershift band was then excised, quantified, and sequenced. The results indicate that there are relatively small differences in binding to templates containing the same p53BS (Fig. 5B). The high-affinity p53BS is bound stronger then the low-affinity across all templates as expected. The control sequences are weakly bound in these experiments 8 to 16 fold below specific binding observed for the sequences containing a p53BS.

**Figure 5.**
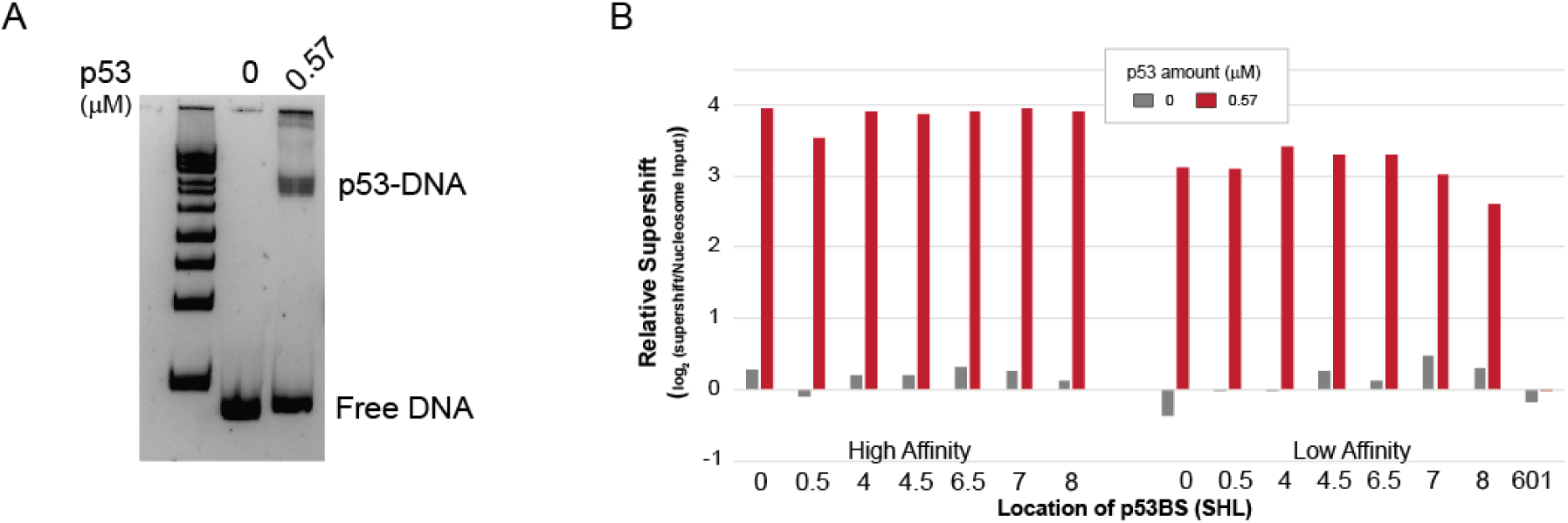
p53 binding to naked DNA. **(A)** 0.5 pmol of DNA containing 16 different p53BS locations were bound to 0.57 µM p53 (4 pmol) **(B)** Relative supershift for each DNA sequence is determined by counting the frequency of each sequence within the supershift and comparing it to non-specific binding to the 601 sequence. This value is then normalized by the input.

### p53 binds nucleosome edges in human lung fibroblasts

To validate the relevance of our *in vitro* binding results we examined p53 induced binding in IMR90 human lung fibroblasts combined with steady state nucleosome occupancy as measured by MNase-seq (Kelly et al. 2012; Sammons et al. 2015). To perform this analysis, we reanalyzed the published MNase-seq and ChIP-seq datasets. The p53 ChIP-seq results are after p53 is induced and defines the specific sites bound by p53. The MNase-seq data was from the same cell line in a steady state without p53 activation and defines the nucleosome position/occupancy before p53 binds. p53 ChIP-seq experiments where performed on IMR90 cells treated with a DMSO control or after p53 activation with nutlin. For each bound site the p53BS was determined from a pre-defined list of p53 motif locations. This step ensures that we are examining direct binding by p53 and have accurate binding locations, but will miss ill-defined or non-consensus binding sequences. The raw MNase-seq data was then extracted, standardized, and visualized centered at the p53BS located within the p53 ChIP-seq peaks. As shown previously average nucleosome occupancy peaks at p53 induced binding sites (Fig. 6A). Closer examination of individual sites after clustering the MNase-seq results shows that p53 TFBS are located within the nucleosome edge flanked by a region of lower nucleosome occupancy (Fig. 6B). These results suggest that p53 induced binding *in vivo* occurs at the edges of nucleosomes, in a manner consistent with our *in vitro* findings.

**Figure 6.**
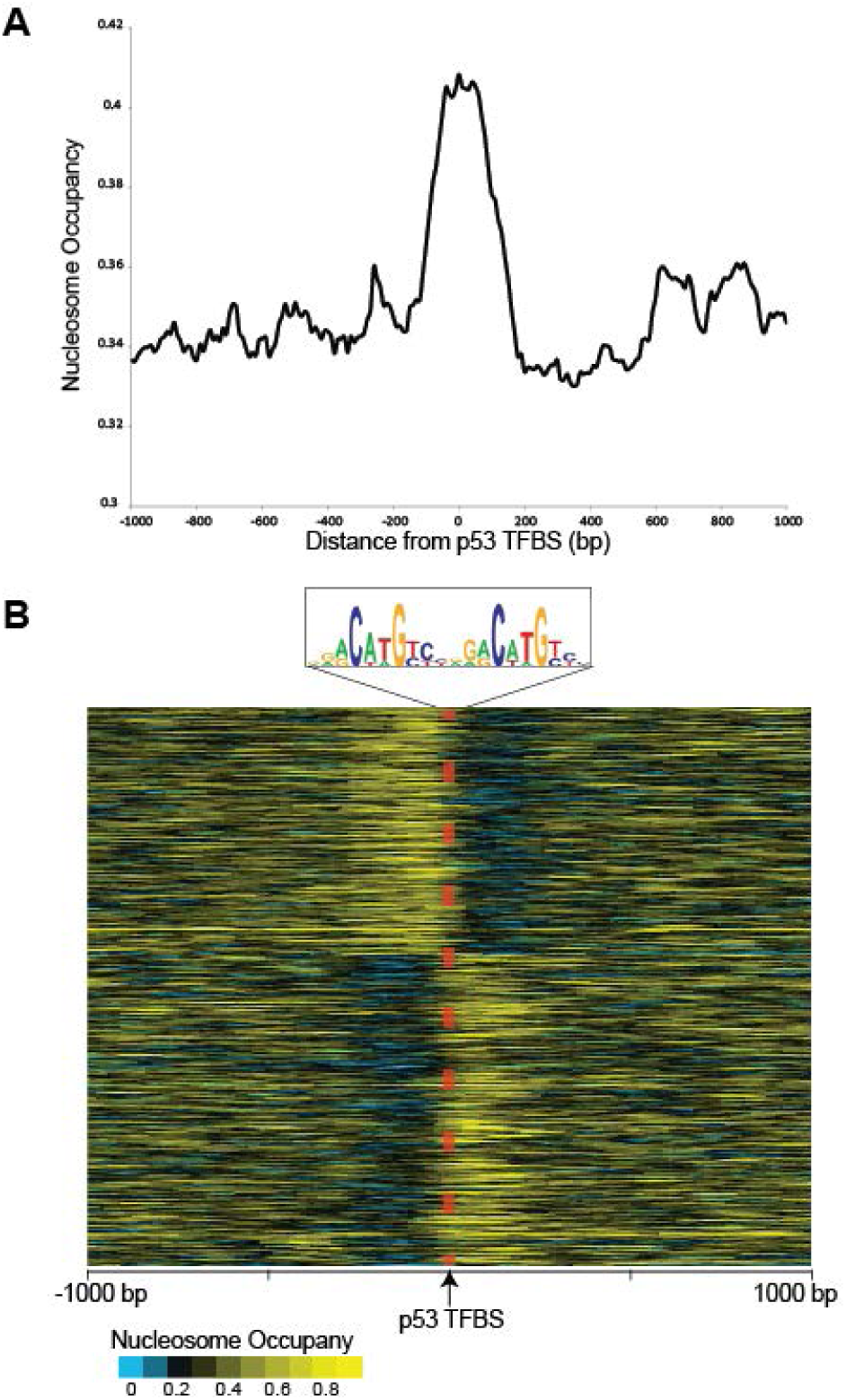
Nucleosome occupancy at p53 nutlin induced binding sites. **(A)** Average nucleosome occupancy at induced p53 binding sites determined from MNase-seq. p53 ChIP-seq results from Sammons et al (2015) with MNase-seq data from Kelly et al. (2012) (Kelly et al. 2012; Sammons et al. 2015). The MNase-seq reads were extracted, standardized (1 billion reads), and extended (120-bp) with ArchTex (Lai et al. 2012). **(B)** Nucleosome occupancy at each p53 TFBS (+/-1000-bp) after nutlin-induced p53 binding. Standardized and extended MNase-seq reads were generated with ArchTex (Lai et al. 2012) and clustered by symmetry (Lai and Buck 2010).

## DISSCUSSION

To identify the rules defining TF targeting in chromatin we developed a new approach to examine the binding characteristics of TFs to nucleosome DNA. Our approach combines traditional proven nucleosome binding assays with next-generation sequencing. This approach allowed the direct comparison of multiple nucleosomes in a single experiment. Each nucleosome template sequence was designed allowing the positioning of high or low-affinity p53BS in various translational and rotational settings. Nucleosomes were then generated from the pool of nucleosome sequences, purified, and competitively bound to p53. Each experiment is controlled at multiple steps. The starting nucleosomes or input to each binding reaction is quantified and sequenced after PAGE (p53-null lane). This input measurement ensures that any variation in nucleosome formation efficiency or starting quantities is corrected for. Within this input lane blank bands corresponding to the supershifted p53-nucleosome complex is also quantified and sequenced. This measurement represents the background for each particular supershift. As can be seen in Fig. 3, for the 0 nM sample (gray bar), there is only slight differences between different nucleosome sequences. In addition to examining the supershifted fragments we also analyzed the nucleosome bands and compared the quantities of each template to the input nucleosomes. In this case we are determining the nucleosome sequences that have shifted out of the nucleosome band after being bound. This analysis does not require us to determine the appropriate supershift band and is resilient to formation of higher order p53-nucleosome complexes seen at higher p53 concentrations. To ensure the accurate absolute quantification of our results we measure the DNA concentration of each gel-excised sample by qPCR. Using this amount with the NGS results we can calculate the % of bound nucleosomes for each nucleosome type across all p53 concentrations. The resulting data were then fitted using non-linear regression to obtain K_D_ values, allowing the examination of both specific and non-specific binding. The methodology we have proposed is not limited to p53-nucleosome binding but can be applied to understand how other pioneer factors bind nucleosomal DNA.

Regardless of the analysis methods we used all approaches consistently showed that TFBS positioning within a nucleosome affect p53-binding capability. p53 displays a strong preference to sites outside the 100-bp surrounding the nucleosome dyad. These results are consistent with a dynamic partial unwrapping near nucleosome edges (Polach and Widom 1995; Li and Widom 2004). In this model, DNA near the entry-exit region is unwrapped from the histone proteins exposing the DNA to TF binding. Once bound the nucleosome can then re-wrap thus kicking off the TF (Luo et al. 2014). Our results for p53 suggest that p53 can access the partially unwrapped nucleosome and remain stably bound. Binding by p53 to nucleosomes differs when compare FoxA which can target is binding site near nucleosome dyads and is likely caused by differences in how the two proteins contact DNA. The canonical pioneer factor FoxA has been shown to be able to bind to target sites located near nucleosome dyad (McPherson et al. 1993; Chaya et al. 2001). FoxA binds nucleosomal DNA on one side as a monomer (Cirillo and Zaret 2007) whereas p53 binds as a tetramer complex that partially wraps around the DNA (Malecka et al. 2009; Emamzadah et al. 2011). These differences in how a TF contacts its binding site may explain the ability of some TFs to bind within the nucleosome core.

Unexpectedly, p53 displays a strong preference to the nucleosome structure itself and binds nucleosomes in a consensus sequence-independent manner with relative high affinity. Our experiments show that control nucleosomes are bound with a K_D_ 38.12 to 38.54 compared with 19.04 for the high-affinity p53BS located in the linker. This two-fold difference in binding affinity is likely an underestimate of the difference between sequence specific and non-specific binding. Examination of the supershift bands at low p53 concentrations show 5 to 11-fold increase of binding to p53BS located in the linker compared to control 601 nucleosomes (Fig. 3). Fluorescence recovery after photobleaching (FRAP) support the model that p53 binds non-specifically to chromatin in the nucleus. Hinow et al (2006) compared wildtype p53 with a DNA-binding mutate and showed indistinguishable nucleus diffusion properties suggesting that p53 had significant sequence-independent binding to chromatin (Hinow et al. 2006). The sequence-independent nucleosome binding is likely due to p53’s CTD. p53 CTD carries a non-sequence-specific DNA-binding activity required for sliding along DNA, which facilitates sequence-specific binding by the DNA binding domain (Sullivan et al. 2018). The sequencing-independent binding observed in our experiments could be caused by either CTD nonspecific binding to the linker or nucleosome DNA.

Our *in vitro* results combined with the *in vivo* binding patterns suggest that p53 is not a FoxA-like canonical pioneer factor; its specific binding capabilities are limited to nucleosome edges flanked by a nucleosome free region. This presents a model for p53 nucleosome binding where p53 encroaches on nucleosomes from an exposed linker region where p53’s CTD binds non-specifically and slides along the DNA. It then gains access to its binding site within the nucleosome edge by the nucleosome partially unwrapping (Fig. 7). Once p53 binds it can recruit histone acetyltransferases further activating the bound enhancer or promoter region (Gu and Roeder 1997; Gu et al. 1997). All members of the p53/p63/p73 family share a similar DNA-binding domain with high sequence and structural homology (Levrero et al. 2000). Structural studies of DNA binding domains for p53, p63, and p73 co-crystallized with DNA target sequences, reveal an overall conserved conformation and DNA-protein contact sites for the three proteins (Chen et al. 2011; Ethayathulla et al. 2012; Ethayathulla et al. 2013). These structural similarities between family members suggest that p63 and p73 will also bind nucleosomal DNA in a similar manner. Genomic studies on p63 have suggested that p63 also acts a pioneer transcription factor during epidermal development by binding at regions of the genome with high encoded nucleosome occupancy (Sethi et al. 2014).

**Figure 7.**
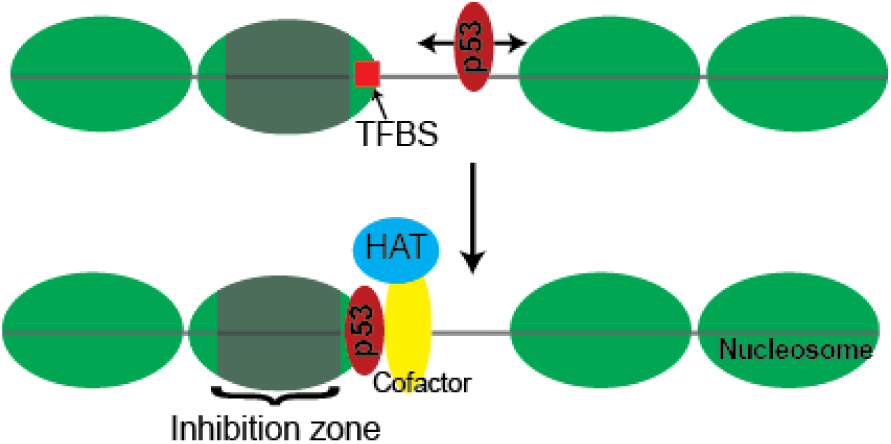
Model of p53 binding to targets within nucleosomes. p53 scans DNA in a sequence-independent manner and target p53BS located at the nucleosome edge. Once bound p53 can recruit cofactors and histone acetyltransferases (HAT).

Chromatin accessibility has been recognized as a pre-requirement for functional activity at regulatory elements (Tsompana and Buck 2014). Our results demonstrate a more meticulous understanding of how positioning within a nucleosome can affect TF binding. In particular, our results demonstrate that positioning within a nucleosome is more important than the affinity of the underlying binding site and small differences in positioning, 50 bp, can dramatically affect TF binding. Therefore, to accurately identify the earliest events during gene activation, high-resolution nucleosome maps will be needed with an understanding of which TFs can target their binding sites within the nucleosome. The methodology we’ve presented here provide a comprehensive approach to examine the rules dictating nucleosome binding by pioneer factors, further studies on all pioneer factors will allow the identification of general themes driving pioneer factor gene regulation.

## MATERIALS AND METHODS

### Design of the nucleosome positioning templates

Nucleosomal templates were derived from the Widom 601 strong nucleosome-positioning sequence (Lowary and Widom 1998; Anderson and Widom 2000). The 601 sequence was first scanned for the presence of sequences similar to the p53 TFBS. The original 601 sequence contains a p53 half-site core (CATG) located just outside the nucleosome edge. To ensure that this sequence doesn’t affect the binding assays we modified this sequence to AGGT. We called it ‘601-modified’, which was regarded as an additional control sequence. The original Widom 601 DNA (601) was still used in the study and had indistinguishable results compare to ‘601-modified’. Moreover, a single half site is not sufficient for p53 binding and has been shown to reduce binding affinity at least 50-fold (el-Deiry et al. 1992; McLure and Lee 1998). p53 TFBS were added at differing positions. Two p53 TFBS were used: an adapted high-affinity ideal sequence: 5’-GGG<u>CATG</u>TCCGGG<u>CATG</u>TCC-3’ (Veprintsev and Fersht 2008; Noureddine et al. 2009) and a natural lower-affinity sequence from the p21 promoter: 5’-AGA<u>CTGG</u>GCATGT<u>CTGG</u>GCA-3’(Westfall et al. 2003). Within each p53 binding sites, there are two cores (underlined above), which are directly bound by p53. CATG and CTGG are the cores in the high-and low-affinity binding sequences, respectively. We design different fragments to ensure that the cores are either in an exposed or concealed orientation as determined by the crystal structure of a nucleosome with the Widom 601 sequence (Makde et al. 2010). Therefore, 14 templates were designed starting from the 217-bp Widom 601 sequence and compared to non-specific binding to 2 control sequences (Table S1).

### *In vitro* nucleosome reconstitution & purification

All 16 synthesized DNA templates were amplified via PCR with the following primers: 5’-GATGGACCCTATACGCGGC-3’, 5’-GGAACACTATCCGACTGGCA-3’. After PCR, all DNA fragments were column-purified (QIAGEN) and quantified. *In vitro* nucleosomes were generated from H2A/H2B dimer and H3.1/H4 tetramer (NEB). All 16 nucleosome sequences were mixed at equal molar amounts. Mixed DNAs were then added to histones at octamer/DNA molar ratios of 1:1 in 2M NaCl. Nucleosomes were reconstituted through salt gradient dialysis as described (Hayes and Lee 1997) which were further purified by 7%-20% sucrose gradient centrifuge (Fang et al. 2016) and concentrated by 50K centrifugal filter units (Millipore, Amicon^R^ Ultra).

Nucleosome formation efficiency was determined by running the reconstituted nucleosomes, before sucrose gradient purification, on a 4% (w/v) native polyacrylamide gel (acrylamide/bisacrylamide, 29:1, w/w, 7 × 10 cm) in 0.5 × Tris Borate-EDTA buffers at 100V at 4 °C. After electrophoresis, the gel was soaked in SYBR Green (LONZA) and incubated at room temperature for 10min, before imaging. DNA was isolated from the nucleosome and free DNA band and sequenced by NGS (see details below). Nucleosome formation efficiency is a measure of the nucleosome formation for each template compared to the standard Widom 601 sequence before nucleosome purification by sucrose gradient (Eq. 1), where *N* is one of the 16 nucleosome sequences, and *601* is the Widom 601 sequence.

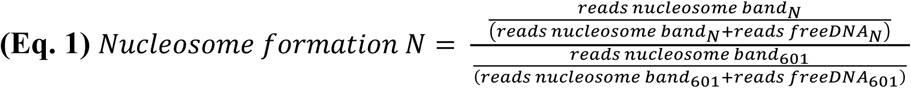

### DNA binding assay followed by EMSA

The protein-nucleosome binding assays were carried out four times with purified nucleosomes mentioned above and human full-length recombinant p53 protein (Abcam cat# ab84768) in 7 µl DNA binding buffer (10mM Tris-Cl, pH7.5; 50mM NaCl; 1mM DTT; 0.25mg/ml BSA; 2mM MgCl^2^; 0.025% P-40; and 5% glycerol), and then incubated for 10 minutes on ice and then for 30 minutes at room temperature. Increasing concentrations of p53 were added to 0.5 pmol purified nucleosomes from p53/nucleosome ratios of 0:1, 0.5:1, 1:1, 2:1 to 4:1. Protein binding was detected by mobility shift assay on 4% (w/v) native polyacrylamide gels (acrylamide/bisacrylamide, 29:1, w/w, 7 × 10 cm) in 0.5 × Tris Borate-EDTA buffers at 100V at 4 °C. After electrophoresis, DNA was imaged by staining with SYBR Green (LONZA).

### DNA isolation and purification

All visual bands were excised from the gel, as well as the bands at the same locations in the other lanes. Each gel slice was processed separately for a total of 80 samples from 4 replicate experiments. In order to extract DNA from the polyacrylamide gel, the chopped gel slices were soaked in diffusion buffer (0.5 M ammonium acetate; 10mM magnesium acetate; 1mM EDTA, pH8.0; 0.1% SDS), and incubated at 50 °C overnight. Supernatant was collected, residual polyacrylamide removed with glass wool, and DNA purified with QIAquick Spin column (QIAGEN). The DNA concentration for each sample was determined by qPCR. Absolute DNA concentration was determined by comparing to a standard curve generated from the control 601 sequence.

### Library construction & sequencing

After we obtained the concentrations of the purified DNA samples, we performed two rounds of PCR to construct sequencing libraries for Illumina sequencing. The first round PCR used forward-amplicon-primer and reverse–amplicon primer (see below). The number of cycles for PCR was determined by the sample concentration determined by qPCR and ranged from 8 to 12 cycles. Each sample was then indexed using Nextera dual indices (Nextera XT Index Primer 1 (N7xx) and Nextera XT Index Primer 2 (S5xx)). After each PCR, reactions were cleaned up with AMPure XP beads (Beckman Coulter). All samples were multiplexed and sequenced in a single lane on the MiSeq using 2 × 150-bp paired-end sequencing. Sequencing and quality control were performed at the University at Buffalo Genomics and Bioinformatics Core.

Forward-amplicon-primer:

5’-TCGTCGGCAGCGTCAGATGTGTATAAGAGACAGGATGGACCCTATACGCGGC -3’

Reverse-amplicon-primer:

5’-GTCTCGTGGGCTCGGAGATGTGTATAAGAGACAGGGAACACTATCCGACTGGCA -3’

### Data analysis

Quality sequence reads were mapped to each specific starting sequence using Blat (Kent 2002). The results were then analyzed relative to control/non-specific binding (relative supershift) or by determining the binding affinity by fitting a binding curve. Relative supershift is determined from the supershift bands and controls technical variability introduced by gel-excision, PCR, NGS-library construction, or NGS sequencing. In this method each specific nucleosome sequence is measured relative to non-specific binding (control 601 fragment).

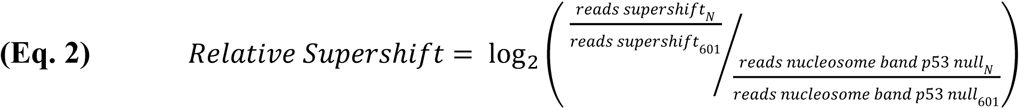

Where *N* is one of the 16 nucleosome sequences, *601* is the control nucleosome sequence, *reads supershift* is the supershift band, *reads nucleosome band* is the nucleosome band in p53-null lane.

Using the DNA concentrations after gel-excision with the number of each nucleosome sequence in a sample the absolute number of a particular nucleosome can be determined for each sample (Eq. 3). Where *N* is one of the 16 nucleosome sequences, *C* is the concentration of the sample after gel extraction (ng/µl), *V* is volume after gel extraction (µl).

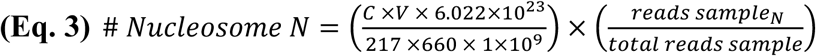

These absolute nucleosome counts when applied to the nucleosome only bands at each p53 concentration are used to determine the % bound nucleosomes for each nucleosome fragment with the following:

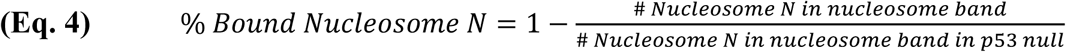

The K_D_ values are then estimated with non-linear regression in Prism (Heffler et al. 2012a).

### Western Blot

Unlike denaturing Western blot analysis, we first performed native PAGE, and then soaked the native polyacrylamide gel in 0.1% SDS for 15 minutes before transfer, and then added SDS to a final concentration of 0.05% in the transfer buffer. After transfer, we incubated the PVDF membrane in 5% acetic acid for 15 minutes to fix the proteins, followed by traditional blocking and antibody incubation steps. Anti-GST antibody (DSHB, P1A12-c) was used to visualize the p53 protein. Anti-Histone H4 polyclonal antibody (Abcam, cat# ab10158) was used to detect the nucleosomes.

### Analysis of p53 induced binding in IMR90

p53 ChIP-seq binding sites identified after nutlin induced binding in IMR90 were obtained from GEO Accession GSE58740. MNase-seq data from proliferating IMR90 were obtained from GEO Accession GSE21823 and aligned to hg19 with Bowtie 2 (Langmead and Salzberg 2012). p53 TFBS from HOMER (Heinz et al. 2010) were intersected to the p53 induced binding sites to define the exact location of binding. p53 ChIP-seq binding sites without a previously defined p53BS were excluded from the analysis. The MNase-seq reads were extracted, standardized (1 billion reads), and extended (120-bp) as done previously with ArchTex (Givens et al. 2012; Lai et al. 2012). Symmetry of resulting MNase-seq dataset at p53BS was then determined with ArchAlign with 0-bp shifts and region reversal enabled (Lai and Buck 2010).

## ACKNOWLEDGEMENTS

We would like to thank Dr. Jeffrey Hayes and Dr. Laxmi Mishra at University of Rochester Medical Center for suggestions of sucrose gradient centrifuge to purify nucleosomes, and UB Genomics and Bioinformatics Core for next-generation sequencing services. This work was partially supported by Mark Diamond Research Fund FA-16-22 to XY and NY State Department of Health C026714 to MJB.

